# Benchmarking five extracellular vesicle proteomics workflows, including low-input Exo-insert and Exo-SP3, for deep mass spectrometry profiling

**DOI:** 10.64898/2026.05.28.728409

**Authors:** Jakub Faktor, Artur Pirog, Anna Biernacka, Ines Papak, Swapnil Bhasvar, Babasaheb Sonwane, Tomasz Marjański, Witold Rzyman, Natalia-Marek Trzonkowska, Sachin Kote

## Abstract

Small extracellular vesicles (sEVs) are key mediators of intercellular communication, influencing diverse pathological processes, including cancer. While mass spectrometry (MS) has enabled the proteomic analysis of sEVs, sample preparation losses remain a critical bottleneck, particularly for scarce tissue-derived sEVs (Ti-EVs). Here, we systematically benchmark five proteomic workflows introducing Exo-insert, a novel single-vessel method, and Exo-SP3, across both Ti-Evs and cell culture-derived sEVs (CCM-EVs) at low input (0.5–4 µg). Exo-insert and Exo-SP3 enable the identification of ∼1100 protein groups from as little as 0.5 µg sEV input. Notably, optimal sample preparation for MS is source-dependent: Exo-insert and Exo-SP3 display divergent performance across sEV sources. Comparative DDA/DIA analyses establish sample preparation as the primary determinant of proteome recovery, offering a practical framework that matches workflows to sEV amounts and source-specific content for biomarker discovery.

## INTRODUCTION

Small extracellular vesicles (sEVs) are lipid bilayer–enclosed particles with diameters of up to 200 nm that are actively secreted by virtually all cell types and play an essential role in intercellular communication. They transport proteins and nucleic acids from donor to recipient cells, modulating cellular function and contributing to physiological and pathological processes [1]. sEVs participate in numerous physiological processes but are also implicated in the development of various pathological conditions, including cancer, neurodegenerative disorders, and autoimmune diseases [2], [3], [4]. Consequently, sEVs have been extensively investigated as potential therapeutic targets, drug delivery vehicles, and diagnostic biomarkers [2], [3]. A particularly important aspect of understanding sEV biology is the characterization of their protein cargo. However, the quantities of sEVs obtained are often insufficient for commonly used mass spectrometry (MS) sample preparation protocols, and depending on the sEV source, signals from true vesicle-associated proteins may be obscured by incompletely removed contaminants, particularly in tissue- or biofluid-derived vesicles. [5]. Reliable analysis therefore, depends on carefully optimized workflows, beginning with exosome isolation and followed by efficient protein extraction and digestion strategies that preserve low-abundance signals. Overcoming issues related to membrane-rich vesicle structure, sample loss, and mass spectrometry compatibility is essential for generating meaningful data. As a result, the development and optimization of sensitive, low-input exosome proteomic pipelines is critical for unlocking the full potential of exosomes as a source of biologically and clinically relevant biomarkers.

Here we benchmark Exo-insert, a novel single-vessel workflow derived from our In-insert method [6], against Exo-SP3 and three conventional sEV proteomics protocols. Exo-insert minimizes handling losses through ultracentrifugation-concentrated, DDM-based sEV lysis in chromatographic inserts, enabling sub-microgram analysis. Exo-SP3, conversely, offers detergent-tolerant bead-based processing with superior robustness.

Across tissue-derived (Ti-EVs) and cell-culture (CCM-EVs) sEVs (0.5–4 µg input), we reveal sEV source-dependent trade-off in processing effectivity. Exo-insert maximizes proteome depth from scarce Ti-EVs, whereas Exo-SP3 enables SDS use. This strong detergent improves protein solubilization and facilitates processing of more complex sample matrices through subsequent bead-based cleanup. This comprehensive comparison provides practical guidance for matching workflows to sEV input amounts and source-specific content, addressing an important bottleneck in clinical sEV proteomics.

## RESULTS

### Characterization of isolated sEVs

For the evaluation of sEV sample MS preparation workflows, we used sEVs isolated by ultracentrifugation (UC) from cell-conditioned culture medium of non-small cell lung cancer (NSCLC) primary cell line (CCM-EVs) or from lung tissue (Ti-EVs). sEVs from both sources were positive for established EV markers (CD9, TSG101, and HSP70) and negative for the common contaminant ApoA1 (**Figure 1 a**). Transmission electron microscopy (TEM) with negative staining revealed the characteristic cup-shaped morphology of the vesicles (**Figure 1 b**), with a median particle size of 145 - 160 nm (**Figure 1 c**).

**Figure 1.**
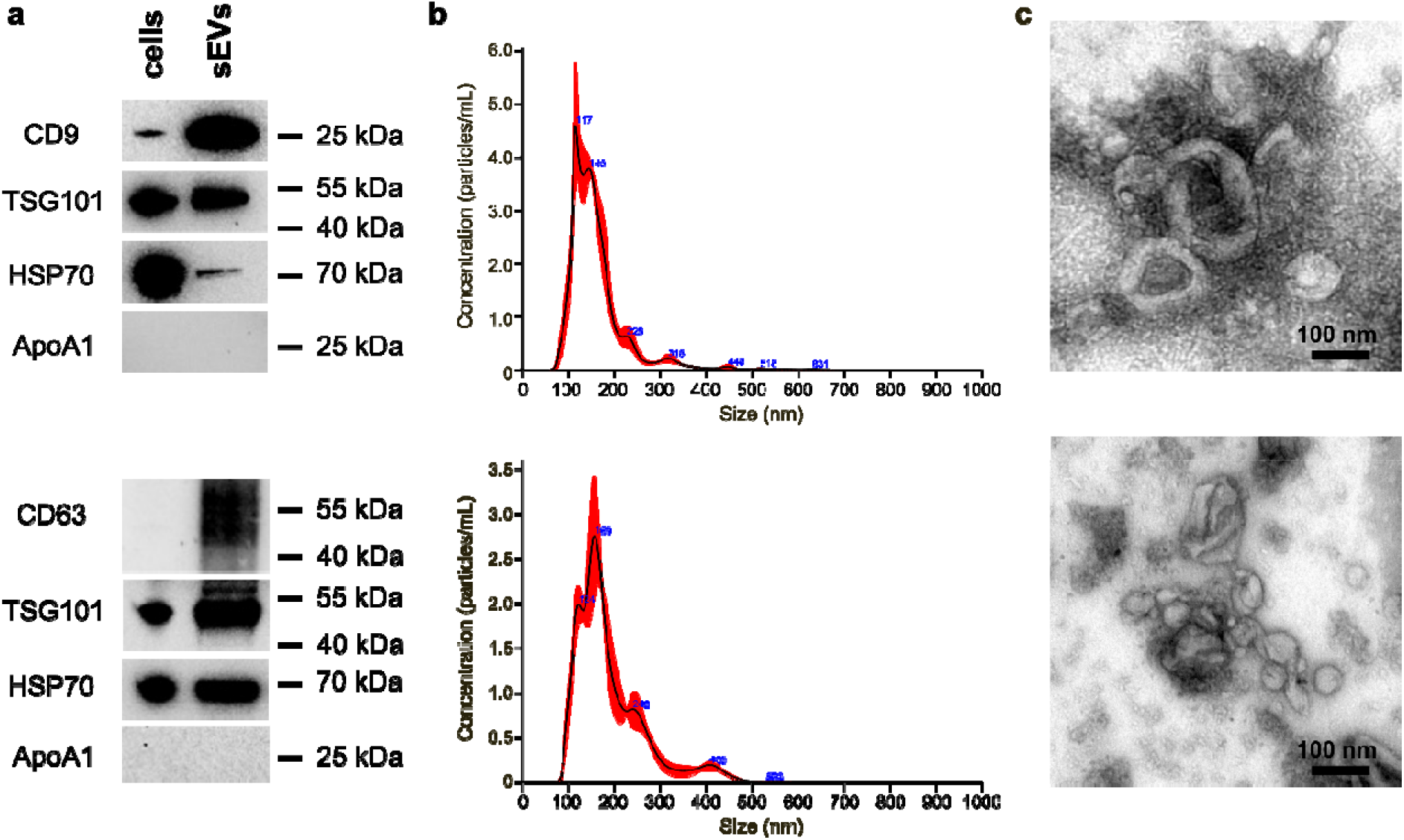
sEV characterization. (a) Western blot analysis of sEV-enriched markers (CD9 or CD63, TSG101, and HSP70) and the common contaminant ApoA1. (b) Size distribution of isolated sEVs, as determined by nanoparticle tracking analysis (NTA). (c) Transmission electron microscopy (TEM) images of isolated particles following uranyl acetate counterstaining. For all panels (a–c), the top images represent Ti- EVs, whereas the bottom images represent CCM-derived EVs.

### Rationale for selection of benchmarked sEV sample preparation methods

Filter-aided sample preparation (FASP), one of the most widely used sEV proteomic sample preparation for MS, requires 5-50 μg of protein input [6], [7], [8], [9], [10], but fails at low amounts of sEVs. Using the FASP protocol with SDS based-lysis (FASP_SDS) we identified only 379 protein groups (90 quantified) from Ti-EV samples, even when 15 μg of material was processed (n=3), with poor peptide separation (**Supplementary table 1, Supplementary figure 1**). This confirms FASP’s known limitations, including on-filter losses and detergent carryover, especially in single-injection, scarce sEV digests. Because FASP_SDS is inadequate for low-input sEV samples, we compared the performance of four workflows representing commonly used and emerging methods (**Table 1**). These included urea-based modification of FASP (FASP_UA) designed to reduce detergent-related interference during downstream mass spectrometry analysis, ammonium bicarbonate (ABC)-based digestion, SP3 bead-based processing, and the Exo-insert approach (single-vessel/in-insert) (**Table 1**). These workflows span lysis chemistry, cleanup complexity, and handling steps, enabling direct assessment of low-input performance.

**Table 1.** Definition of sEV sample-preparation workflows for mass spectrometry. The table summarizes lysis buffer composition, processing format, hands-on time, number of pre-digestion steps, desalting needs, and sEV input requirements (Exo-insert, ABC, Exo-SP3, FASP_UA, and FASP_SDS). It also indicates sample input state (pellet vs. solution), tolerance to detergents/urea, and key advantages and limitations, providing a concise basis for comparison.

| Principle | Direct/in-solution (no desalting) | Direct/in-solution (with desalting) | On-bead (SP3-based) | On-filter |  |
| --- | --- | --- | --- | --- | --- |
| Method name | 1. Exo-insert | 2. ABC | 3. Exo-SP3 | 4. FASP-UA | 5. FASP-SDS |
| Origin | Modified from Faktor et al. [11] | Rao et al. [12] | Modified from Hughes et al. [13], Muraoka- et al. [14], Cross et al. [15] | Modified from Wisniewski et al. [16], Patel et al. [17] , Ding et al. [18], Wang et al. [19] | Modification of Wisniewski et al. [16], Wu et al. [10] |
| Lysis buffer | 0.2% (w/v) DDM and 10 mM DTT | 50 mM NH<sub>4</sub> HCO<sub>3</sub> | 2% SDS (w/v), 0.1 M Tris/HCl, 0.1 M DTT, pH 8.5 | 8 M urea, 0.1 M Tris/HCl, pH 8.5 | 4% (w/v) SDS, 0.1 M Tris/HCl pH 7.6, and 0.1 M DTT |
| Processing time up to digestion | ~ 30 min | ~ 30 min | ~ 6 h | ~ 6 h | ~ 6 h |
| N steps | 4 | 2 | 8 Artur confirm | 9 | 9 |
| Desalting | NO | YES | YES | YES | YES |
| Format | In single container, direct | Direct | On-beads | On-filter | On-filter |
| Required sample status/format | PBS washed sEV pellet with minimal MS-incompatible substances | PBS washed sEV pellet with minimal MS-incompatible substances | sEVs in solution; high SDS concentration compatible | sEVs in solution; high urea concentration compatible | sEVs in solution; high SDS concentration |
| Main limitations | Requires ultracentrifuged pellet; low-volume handling; sensitive to residual contaminants | Sample loss during desalting; dilution | Multiple handling steps may increase sample loss | Multiple handling steps may increase sample loss | Multiple handling steps may increase sample loss; residual SDS may affect LC-MS if not sufficiently removed |
| Main advantages | Rapid workflow; minimal surface contact; low consumable demand | Rapid workflow; low handling complexity | Compatible with diverse lysis buffers and MS incompatible detergents | Reduced sensitivity to residual contaminants; compatible with urea lysis | Reduced sensitivity to residual contaminants |
| Scalability | Adaptable to 96 well/insert format after pellet transfer | Adaptable to 96 well/insert format after pellet transfer | Limited parallelization | Limited parallelization | Limited parallelization |
| Risk of sample loss | During the pellet transfer | During the pellet transfer and desalting | Increased due to repeated surface contact | Increased due to repeated surface contact | Increased due to repeated surface contact |
| Contamination risk | Higher dependence on initial sample purity | Moderate | Low | Low | Moderate to high if SDS removal is incomplete |
| Reported usage | First application in this study | [12] | [14], [15], [20], [21] | No reports for sEV lysis and processing via urea-FASP | [6], [7], [8], [9], [10] |

### Novel aspects of emerging Exo-insert proteomics method processing sEVs

The Exo-insert sample preparation workflow is shown in **Figure 2**. The Exo-insert differs from canonical workflows and all other methods listed in **Table 1** by enabling effective proteomic processing of sEVs within a single chromatographic insert. Compared to the previous In-insert method [11], the Exo-insert workflow incorporates modified lysis and protein extraction conditions along with a reduced sample volume, allowing the entire sEV input to be processed and analyzed in a single LC-MS/MS injection using a conventionally configured nano-LC system. The workflow combines concentration and washing of the isolated sEV sample by ultracentrifugation, transfer of the sample to a minimal volume in an insert tube, and direct processing in the same insert. N-dodecyl β-D-maltoside (DDM) based lysis is employed to enable efficient processing within the single insert. This design reduces handling steps and limits sample exposure to surfaces or/and purification procedures. As a result, the method operates at a reduced processing volume, with fewer transfer steps, and is compatible with parallelized handling formats, including 96-position insert plates.

**Figure 2.**
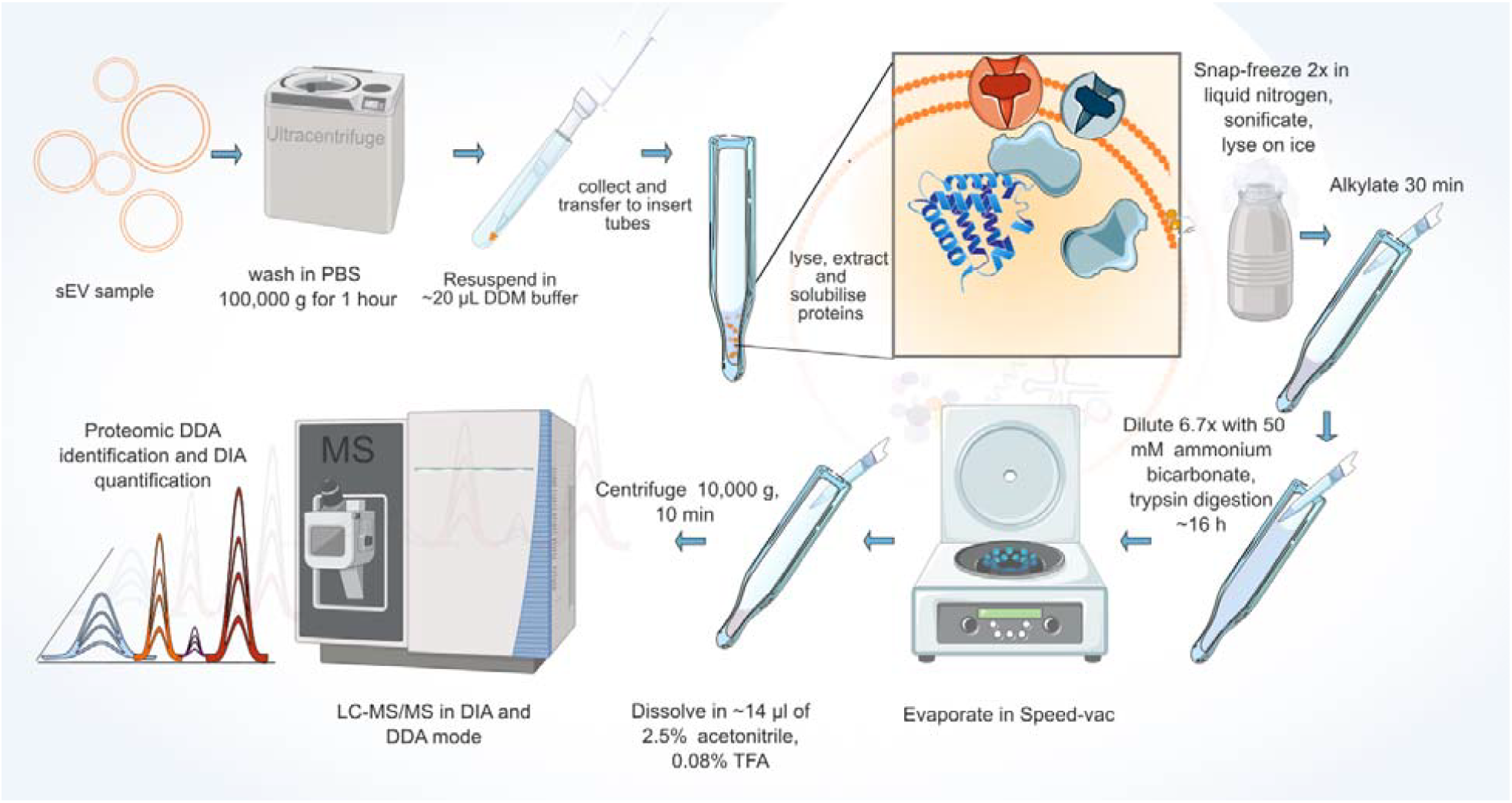
Optimized Exo-insert proteomics workflow for processing sEVs for direct bottom-up proteomics within a chromatographic insert. The sample is processed at the microliter scale in a single container insert, minimizing losses associated with surface adsorption while maintaining the sample concentrated and free of mass spectrometry-incompatible substances, providing an efficient proteomic sample preparation platform. The Implementation of both data-dependent (DDA) and data-independent (DIA) LC-MS/MS workflows enabled sub-microgram sEV input sensitivity while maintaining insight into quantitative proteomic signatures.

### Comparative evaluation of five sEV MS sample preparation methods using CCM-EVs and Ti-EVs

To compare the performance of the four protocols described in **Table 1** (excluding FASP-SDS), we used CCM-EVs, which could be produced in higher quantities, enabling side-by-side comparisons of various workflows. Samples were prepared in triplicate and analyzed across multiple input amounts (0.5–4 µg), except for 2 µg of CCM-EVs processed by FASP_UA, which was not included in the final analysis.

Each resulting tryptic peptide digest was split and analyzed by DDA and DIA acquisition on an Exploris 480 MS to reveal the contribution of the MS acquisition strategy to protein identification and quantification. DDA data performance was assessed by counting identified protein groups (**Figure 3 a**) and peptides (**Figure 3 b**), while DIA performance was evaluated by counting quantified protein groups and peptides with valid quantitative values (**Figure 3 c, d**).

**Figure 3.**
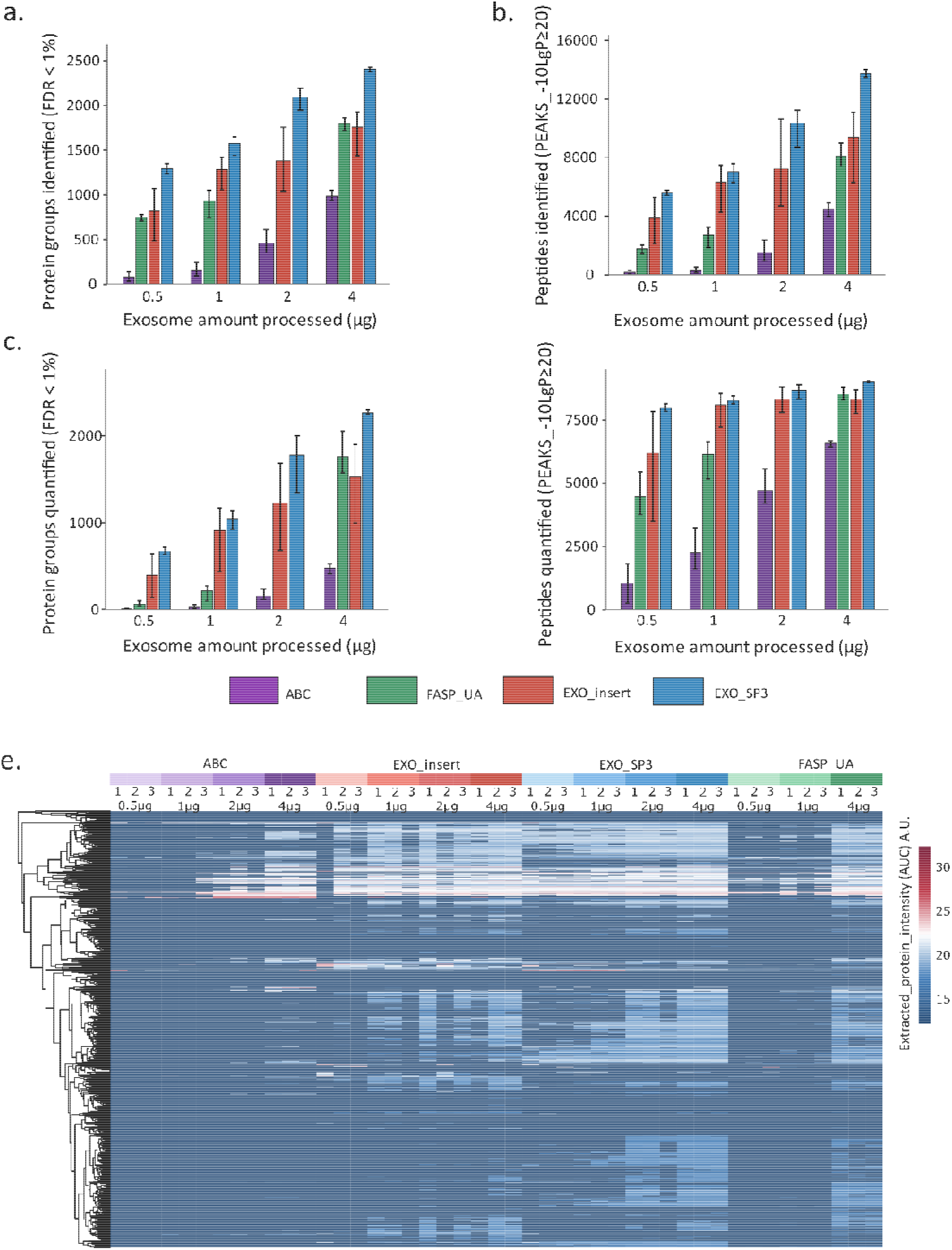
Comparison of CCM-EV proteome coverage across preparation methods, input amounts, and acquisition strategies. CCM-EVs were processed using four preparation methods (ABC, Exo-insert, Exo-SP3, FASP_UA) across multiple sEV input amounts (0.5-4 µg, except 2 µg for FASP_UA), each analyzed in technical triplicate. Peptide samples were split for analysis by DDA and DIA on an Exploris 480 platform, and datasets were processed separately. (a) Stacked bar plots showing the mean number of protein groups identified by DDA. (b) Stacked bar plots showing the mean number of peptides identified by DDA. (c) Stacked bar plots showing the mean number of protein groups quantified by DIA. (d) Stacked bar plots showing the mean number of peptides quantified by DIA. Error bars in bar plots (a-d) show minimum and maximum values. (e) Heatmap of DIA-derived protein intensities across all method replicates.

Exo-SP3 yielded the broadest proteome coverage (**Figure 3 a - e**) and the highest number of protein groups identified and quantified (**Figure 3 a, c**), followed by Exo-insert and FASP_UA, which likely benefits from detergent-free urea lysis and snap-freezing. ABC performed the worst.

Comparison of the number of protein groups identified (**Figure 3 a**) and quantified (**Figure 3 c**) revealed systematic differences between orthogonal DDA and DIA measurements. At low sEV input, DIA quantification sensitivity drops more rapidly than DDA identification, most notably at 0.5 µg, where the discrepancy between identified and quantified protein groups increased to over nine-fold on average across methods and was inversely proportional to the expected sensitivity of the method (**Supplementary table 2**).

**Table 2.** Overall qualitative rating (1 = poor, 5 = high) of the 5 methods by integrating experimental insights from **Figure 3-5** with key sample preparation parameters from **Table 1**, evaluating the balance between protocol complexity, robustness, and proteome depth. The overall assesment performed by the providing possibilities of clinical biomaker discovery.

| Method | Simplicity | Speed | Proteome Coverage | Sample Loss Risk | Robustness (dirty samples) | Throughput | Overall | Comment |
| --- | --- | --- | --- | --- | --- | --- | --- | --- |
| Exo-insert | 5 | 5 | 4-5 | 4 | 2 | 5 | 4.4 | Best balance; strongest for low input & throughput |
| Exo-SP3 | 2 | 3 | 4-5 | 3 | 5 | 2 | 4.2 | Best overall for depth & robustness |
| ABC | 5 | 5 | 2 | 4 | 2 | 5 | 3.7 | Fast but limited depth |
| FASP_UA | 3 | 3 | 4 | 3 | 4 | 2 | 3.7 | Solid, reliable middle ground |
| FASP_SDS | 2 | 3 | 1 | 4 | 4 | 2 | 3.3 | Penalized by low coverage despite robustness |

However, trends observed at the protein level were only partially consistent with those at the peptide level (**Figure 3 c, d, Supplementary table 2**). In DIA, the number of quantified peptides (**Figure 3 d**) increased at low sEV input amounts but quickly reached saturation, with a plateau between 1 and 2 µg sEVs processed. In contrast, quantitated protein group counts in DIA continue to rise across the full range of input amounts (**Figure 3 c**), despite the early stabilization at the peptide level (**Figure 3 d**). On the other hand, for DDA, both peptide and protein group identifications increase progressively with increasing sample load, exhibiting a more linear relationship (**Figure 3 a, b**), highlighting differences in PEAKS algorithms for proteomic quantification and identification.

Our results in **Figure 3 d** indicate that Exo-SP3 and Exo-insert provide the most effective sampling in DIA at low sEV input, likely reflecting the most efficient protein extraction and correspondingly lower limits of detection (LOD) and quantification (LOQ) compared to the other methods included in the comparison (**Supplementary figure 2, Figure 3 a-e**). These methods exhibit relatively small differences between identified and quantified protein groups (2-fold and 2.5-fold, respectively) (**Supplementary table 2, Figure 3**). At higher sEV loadings the method-dependent differences in DIA sampling diminish (**Figure 3 a, c, d**) with the exception of ABC method. While the average discrepancy between identified and quantified protein groups decreases to ∼2.2-fold and then to ∼1.5-fold across all compared methods, indicating near convergence between identification and quantification depth under these conditions (**Supplementary table 2, Figure 3 d**).

Overall, Exo-SP3 and Exo-insert emerged as the most effective and sensitive methods in our dataset. Moreover, the results indicated that Exo-insert and Exo-SP3 exhibit broadly comparable performance in the full method comparison, including detection limits, number of identified peptides and proteins, and overall signal intensity. However, these differences appear relatively moderate compared with those of other methods (FASP_UA), suggesting that a direct head-to-head comparison may reveal more pronounced performance differences. Accordingly, these two methods were selected for a more detailed comparative analysis.

### In-depth comparative evaluation of Exo-SP3 and Exo-insert workflows for sEV proteomic sample preparation

To further evaluate the performance of Exo-SP3 and Exo-insert across different sEV sources, we analyzed scarce Ti-EVs alongside CCM-EVs to assess source-specific benchmark performance. Input amounts ranged, as previously applied, from 0.5 to 4 µg of sEV material. Surprisingly, for Ti-EVs, Exo-insert consistently outperformed Exo-SP3 (and all other methods, as shown in **Supplementary figure 2**) across all input amounts, unlike in CCM-EVs (**Figure 3, Figure 4**). At 0.5 µg input, Exo-insert enabled identification of 1462 protein groups (-10logP > 20) in DDA (**Figure 4 a**), and quantification of 867 protein groups in DIA (**Figure 4 b**), increasing to 1741 protein groups identified, and 1458 protein groups quantified at 4 µg Ti-EVs input. Additionally, Exo-insert displayed relatively high protein intensities in the DIA heatmap even at 0.5 µg Ti-EVs input, revealing distinct clusters with uniquely elevated protein signals, suggesting reduced method extraction bias (**Figure 4 e, Supplementary figure 2**). Exo-SP3 consistently showed slightly lower performance, identifying 796-1490 protein groups and quantifying 274-1105 protein groups across 0.5 - 4 µg Ti-EVs Figure 4 c-d, whereas in CCM-EVs it outperformed the Exo-insert (**Figure 3, Figure 4**). However, heatmap analysis showed more balanced selectivity and extraction efficiency for CCM-EVs than for Ti-EVs, regardless of whether Exo-insert or Exo-SP3 was used. In addition, CCM-EV exhibited reduced method-specific differences in protein clusters compared with Ti-EVs (**Figure 4 e**).

**Figure 4.**
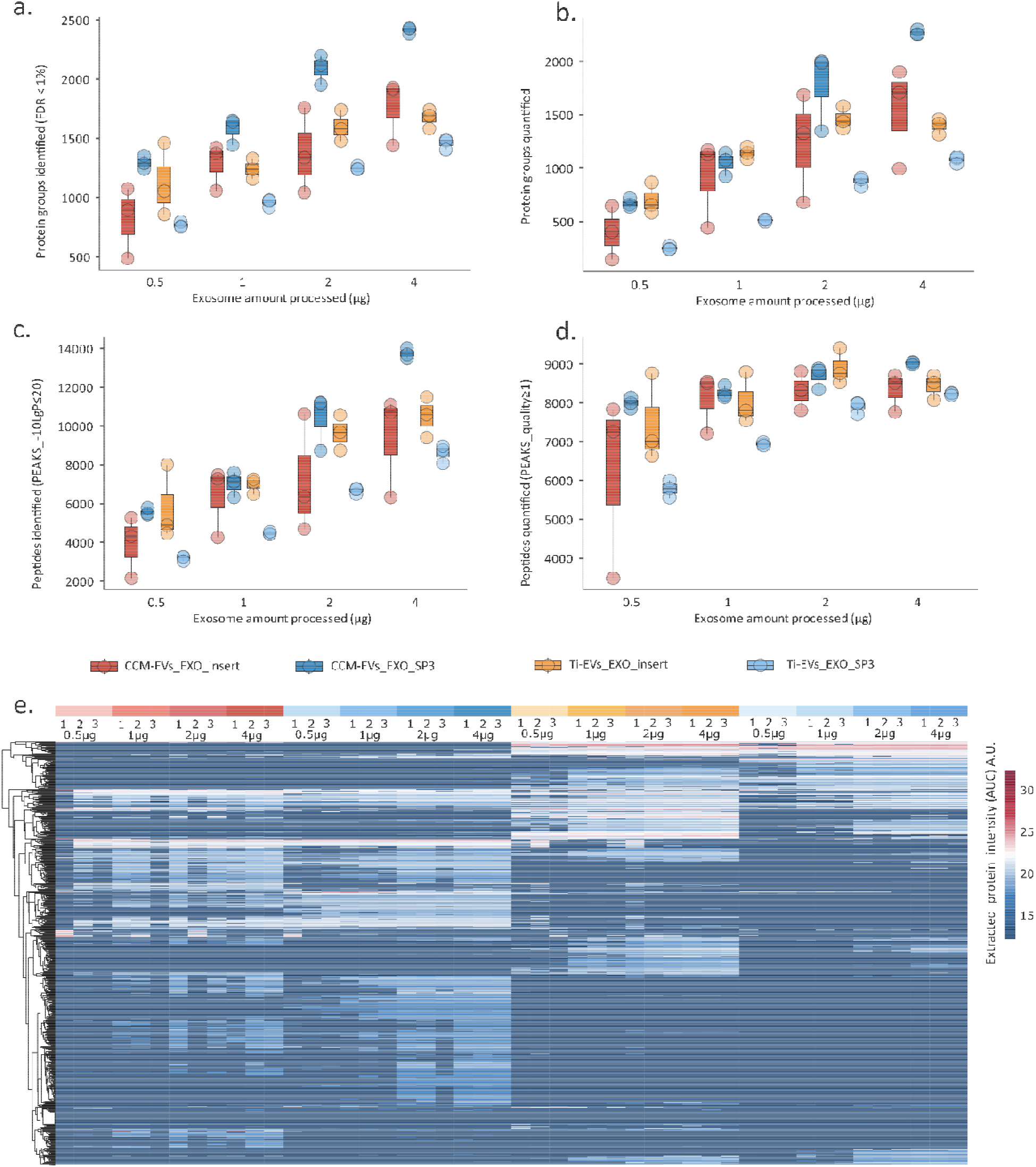
Performance, variability and selectivity of Exo-SP3 and Exo-insert across sEV inputs (0.5, 1, 2, 4 µg). Side-by-side box plots show triplicate variance for Ti-EVs and CCM-EVs: (a) protein groups identified (DDA), (b) peptides identified (DDA), (c) quantified protein groups (DIA), and (d) quantified peptides (DIA). Boxplots display median, interquartile range, and min–max whiskers. (e) Heatmap of DIA-derived protein intensities across all conditions reveals distinct protein intensity patterns, highlighting method-specific protein clusters driven by differences in protocol steps, selectivity, and extraction efficiency.

Exo-SP3 had lower variability in the number of identified and quantified protein groups and peptides, whereas Exo-insert exhibited higher variability across all input amounts and sEV sources (**Figure 4 a-d**). For both methods, the variability was higher in CCM-EVs than in Ti-EVs.

The increased replicate variability observed for Exo-insert is likely attributable to the operator-dependent steps of pellet dissolution and material transfer from the UC tube to the insert tube. While Exo-insert consistently shows higher sensitivity in Ti-EVs, Exo-SP3 outperforms it in CCM-EVs, both in terms of reproducibility and overall protein yield (**Figure 4, Supplementary figure 2**).

We further evaluated reproducibility, variability and differences among proteotypes from our dataset using PCA, UMAP and Pearson correlation heatmap for both methods and both sEVs sources at different input amounts (**Figure 5 a-c**). PCA (**Figure 5 a**) showed a clear separation along principal component 1 (PC1), indicating that sEV source was the most dominant source of the variability, followed by processing method. The second principal component (PC2) in combination with PC1, further resolved the samples according to sEV input across the compared methods. In parallel, UMAP analysis (**Figure 5 b**) revealed 4 distinct clusters, each corresponding to a unique combination of processing method and sEV source. The only exception was 0.5 µg Ti-EV input processed via Exo-insert, which formed a separate outlier cluster.

**Figure 5.**
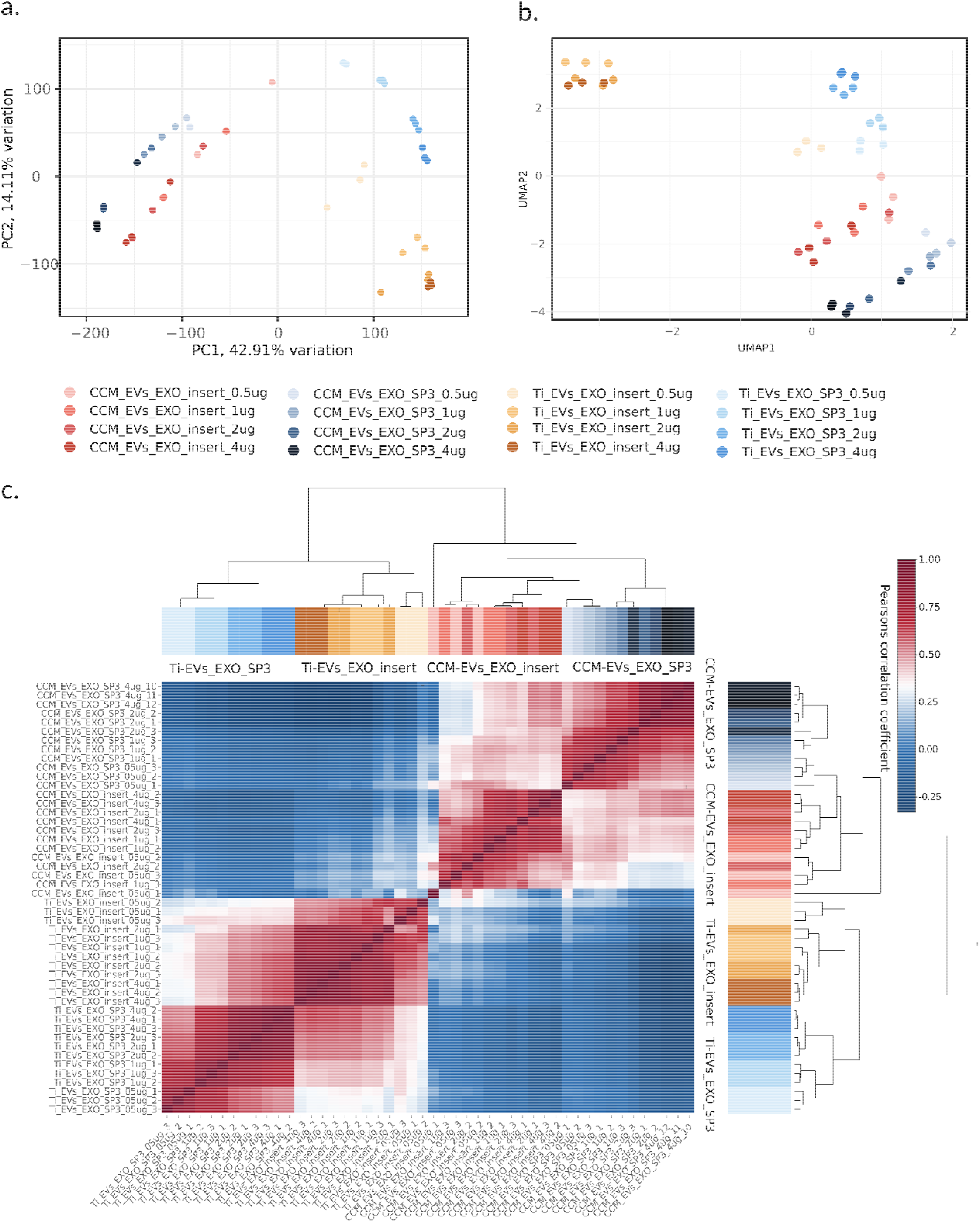
Variability across DIA replicates for Exo-insert and Exo-SP3 methods at 0.5, 1, 2, and 4 µg loadings of Ti-EVs and CCM-EVs, assessed by (a) PCA, (b) UMAP, and (c) Pearson correlation heatmap. Exo-insert shows greater variability than Exo-SP3 across both sample types. Inter-method correlations suggest distinct, method-dependent proteome profiles. Variability is higher in CCM-EVs than in Ti-EVs for both methods.

As expected, the Pearson correlation heatmap of sEV proteotypes (**Figure 5 c**) revealed the highest correlations between replicates (equal sEV input amounts from the same source processed and analyzed using the same method). Additionally, it further refined our previous observations and confirmed that the primary source of variability among the samples was sEV source, followed by the processing method. Notably, the lower inter-replicate correlation coefficients for Exo-insert compared to Exo-SP3 are consistent with the observed higher variability in number of identified peptides and protein groups in Exo-insert samples (**Figure 4 a-d**). Across technical replicates, Exo-insert showed Pearson correlation coefficients ranging from r>0.65 (0.5 µg) up to 0.95 (4 µg) for Ti-EVs, and from r>0.34 (0.5 µg) up to 0.88 (4 µg) for CCM-EVs, with a clear increase in correlation at higher sEV input amounts. In contrast, Exo-SP3 demonstrated higher reproducibility, with correlations ranging from r>0.92 (0.5 µg) up to 0.97 (4 µg) for Ti-EVs, and from r>0.69 (0.5 µg) up to 0.95 (4 µg) for CCM-EVs.

Inter-method correlation coefficients for matched sEV sources ranged from 0.25 to 0.65 (Ti-EVs) and -0.06 to 0.56 (CCM-EVs) (**Figure 5 c**). Collectively, these data suggest improved reproducibility of Exo-SP3 relative to Exo-insert method, while lower inter-method correlations indicate complementary method performance, reflecting differences in selectivity and extraction efficiencies.

### Summary of key sEVs proteomic sample preparation parameters and performance metrics

To complement the experimental results shown in **Figure 3-5**, we summarized key sample preparation parameters and performance metrics, including protocol complexity, robustness, and proteome depth (**Table 2**). Notably, SDS-based FASP_SDS applied to Ti-EVs is consistently inferior to urea-based FASP_UA on CCM-EVs, highlighting the general advantage of detergent-free conditions at low sEV input rather than a bias related to sEV origin (**Supplementary figure 2**). Nevertheless, overall, Exo-SP3 and Exo-insert consistently emerged as the most effective and sensitive methods across sample sources, while ABC performed worst. In addition, the results indicated that Exo-insert and Exo-SP3 exhibit broadly comparable performance in the full method comparison, including detection limits, number of identified peptides and proteins, and overall signal intensity.

## DISCUSSION

sEVs carry nucleic acids, metabolites, and proteins that mirror the physiological or pathological state of their tissue of origin [22], [23], positioning them as rich sources for biomarker discovery [24], [25], [26]. Yet, their proteome analysis from scarce material, particularly Ti-EVs [27], remains constrained less by MS sensitivity than by sample preparation losses. Here, we directly benchmark four workflows across Ti-EVs and CCM-EVs, revealing that optimal performance is source-dependent: Exo-insert maximizes depth in Ti-EVs, while Exo-SP3 provides superior robustness in CCM-EVs.

Although sEV proteomics workflows have been previously explored [28], [29], direct benchmarking under low-input conditions remains limited [6], despite ongoing efforts to develop more sensitive approaches [6], [15], [30]. Commonly used sEV proteomics workflows, including FASP [6], [7], [8], [9], [10], [31], in-gel digestion [32], [33], [34], [35], and in-solution digestion [36], [37], [38], often perform sub-optimally at low input due to surface adsorption, detergent MS incompatibility, multi-step handling, or poor recovery of tryptic peptides [39], [40], [41]. This gap is particularly relevant for Ti-EVs, where restricted tissue availability limits sEV yields while systematic, input-resolved comparisons across biologically distinct sEV sources are lacking. To address this gap, we evaluated SDS-based FASP (FASP_SDS), urea-modified FASP (FASP_UA), direct digestion in ammonium bicarbonate (ABC), SP3 (Exo-SP3), and our novel Exo-insert workflow across 0.5–4 µg sEV inputs (15 µg for FASP) prepared in triplicate, using both DDA and DIA.

FASP_SDS performed poorly, identifying only 379 protein groups (90 quantified) even at 15 µg Ti-EVs, with suboptimal peptide separation, confirming its unsuitability for low-input sEVs. FASP_UA improved this (>1800 proteins at 4 µg CCM-EVs), slightly exceeding previously reported FASP-based EV datasets (1200 protein groups at 5 µg EVs) [6], while ABC lagged, consistent with previously reported low-input in-solution approaches, likely reflecting inefficient ammonium bicarbonate lysis and cumulative losses during mandatory C18 cleanup [6], [12]. Exo-SP3 and Exo-insert dominated, but source-specifically. At 0.5 µg Ti-EVs Exo-insert (identified 1462 and quantified 867 protein groups) outperformed Exo-SP3 (identified 796 and quantified 274 protein groups). Conversely, for CCM-EVs, Exo-SP3 showed the best performance (identified 1354 and quantified 719 protein groups at 0.5 µg sEV input), followed by Exo-insert and FASP_UA. This likely reflects higher tolerance of Exo-SP3 to MS-incompatible molecules, whereas residual matrix effects from varying components of the sEV-associated material (sEV corona) or trace media contaminants may reduce the effectiveness of MS analysis in Exo-Insert. Notably, a previous SP3-based workflow reported ∼1,900 protein groups at 0.5 µg of colon carcinoma-derived sEVs with similar inter-replicate variability patterns [6], while no equivalent benchmark exists for Exo-insert.

Novel Exo-insert addresses preparation bottlenecks such as contact with plastic surfaces, desalting media, and molecular cut-off membranes through single-vessel processing in a chromatographic insert (∼20 µl), using MS-compatible DDM lysis post-ultracentrifugation, minimizing transfers and surface exposure. Unlike prior insert methods, it enables full-input analysis via conventional nano-LC, reducing dilution and loss while enabling scaling to 96 insert position plates. In addition, heatmaps revealed Exo-insert’s reduced extraction bias in Ti-EVs, with distinct high-intensity protein clusters. This also likely reflects detergent chemistry, DDM preserves hydrophobic structure and cleavage accessibility, whereas SDS used in SP3 and FASP_SDS promotes more aggressive unfolding, which rather enhances extraction of resistant protein complexes [42], [43], [44]. Of note, conventional sEV lysis typically relies on strong MS-incompatible detergents such as SDS [29], [45], [46], NP-40 [47], Triton X-100 [8], or RIPA [35], which require removal steps that introduce losses, steps that are not necessary with Exo-insert.

Across input titrations processed through Exo-SP3 and Exo-insert, DDA showed linear increases in identifications with input, while DIA exhibited peptide saturation at 1–2 µg, yet continued protein-group gains via refined inference likely driven by improved peptide spectral evidence. Exo-SP3 and Exo-insert achieved the tightest identification-quantification gaps (∼2-fold at low input), together indicating superior extraction efficiency and lower LOD/LOQ (**Supplementary table 2**). This behavior was not observed for lower-performing workflows such as ABC, whose peptide level saturation likely lies beyond 4 µg sEV input, perhaps due to lesser efficiency of ammonium bicarbonate lysis and adsorption related peptide losses.

Multivariate analyses confirmed sEV source as the dominant variance driver (PC1 in PCA and UMAP), followed by method, with sEV input contributing the least (**Figure 5 a–c**). Exo-SP3 showed higher reproducibility (r > 0.92–0.97) than Exo-insert (r > 0.65–0.95), especially in CCM-EVs, while inter-method correlations were generally low (0.25–0.65), underscoring method complementarity (**Figure 5 c**). These source-method interactions suggest preparation selectivity: DDM in Exo-insert favors hydrophobic accessibility in relatively clean Ti-EVs, whereas Exo-SP3 enables the use of SDS, a strong detergent that improves protein solubilization and facilitates processing of more complex residual sEV material related matrices through subsequent bead-based cleanup [42], [43], [44].

In summary, low-input sEV proteomics represents a source- and input-dependent optimization challenge in which selecting a sample preparation strategy appropriate for a given vesicle source and input amount has a greater impact on proteome recovery than column loading or MS acquisition mode. Exo-insert provides deeper proteome coverage for Ti-EVs and may therefore be particularly suitable for biomarker-oriented applications, while Exo-SP3 ensures greater robustness across different sEv material-related matrices. Altogether, our results provide researchers with a decision framework to extract maximal biological insight from scarce material (**Table 2**).

## MATERIALS AND METHODS

### Samples

Tumor specimens were obtained via surgical resection from non-small cell lung cancer (NSCLC) patients prior the chemotherapeutic treatment. Corresponding samples of healthy lung tissue were collected in parallel. Samples were obtained from the Department of Thoracic Surgery, Medical University of Gdańsk. The study was conducted with ethical approval from the Independent Bioethics Committee for Scientific Research at the Medical University of Gdańsk (approval no. NKBBN/880-329/2016).

### Tissue digestion and primary cell line establishment

Tumour tissues were mechanically fragmented and enzymatically digested using a Tissue Digestion Kit (Real Research), supplemented with DNase I (40 U/mL; GoldBio), for 45 min at 37°C. The digested material was subsequently passed through a 70 µm cell strainer and washed with PBS. The resulting cell suspension was centrifuged at 500 × g for 5 min. The tissue supernatant (TSN) was retained for small extracellular vesicle (sEV) isolation, whereas the cell pellet was used for the establishment of primary cell lines.

Cells were subjected to immune cell depletion using a CD45^+^ depletion kit (StemCell Technologies) and subsequently cultured in TumorPlus medium (Capricorn Scientific) supplemented with penicillin (100 U/mL) and streptomycin (100 µg/mL) (Gibco) at 37 °C in a humidified atmosphere containing 5% CO_2_.

For sEV isolation, the established primary lung cancer cell line was cultured in Dulbecco’s Modified Eagle’s Medium (DMEM) supplemented with 10% EV-depleted fetal bovine serum (FBS) (Gibco), penicillin (100 U/mL), and streptomycin (100 µg/mL) (Gibco) for 72 h. Conditioned cell culture medium (CCM) was then collected for sEV isolation. EV-depleted FBS was prepared by ultracentrifugation at 100,000 × g for 19 h.

### sEV isolation by ultracentrifugation

To isolate sEVs, CCM or TSN was processed by differential centrifugation. Samples were first centrifuged at 300 × g for 10 min to remove cells and large debris, followed by centrifugation at 2,000 × g for 20 min to eliminate insoluble proteins and apoptotic bodies. The supernatant was then centrifuged at 11,000 × g (average) for 30 min using an F37L-8×100 rotor (Sorvall WX+ ultracentrifuge, Thermo Fisher Scientific) to remove microvesicles.

Subsequently, the resulting supernatant was filtered through a 0.2 µm filter and ultracentrifuged at 100,000 × g (average) for 123 min to pellet sEVs. The pellet was resuspended in particle-free PBS and centrifuged again under the same conditions to remove residual medium components. All centrifugation steps were performed at 4°C. The final sEV pellet was resuspended in particle-free PBS and stored at −80°C until further use.

### sEV characterization

sEV concentration and size distribution were determined by nanoparticle tracking analysis (NTA) using a NanoSight NS300 instrument (Malvern Panalytical) equipped with a 488 nm laser. For each sample, five 60 sec recordings were acquired. Protein concentrations were measured using the Qubit Protein Assay Kit and a Qubit 4 Fluorometer (Thermo Fisher Scientific).

Transmission electron microscopy (TEM) of sEVs was performed at the University of Gdańsk (Laboratory of Electron Microscopy) as a paid service. Samples were transferred onto carbon film-coated, 300-mesh copper grids (Agar Scientific) and subjected to negative staining with 1.5% uranyl acetate (BD Chemicals Ltd.). Images were acquired using a Tecnai G2 Spirit BioTWIN transmission electron microscope (FEI Inc.).

For Western blot analysis, cell and sEV samples were lysed in RIPA buffer supplemented with cOmplete Mini Protease Inhibitor Cocktail (Roche) for 15–30 min on ice, followed by mixing with 5× Laemmli sample buffer (60 mM Tris-HCl, pH 6.8; 2% SDS; 10% glycerol; 5% β-mercaptoethanol; 0.2% bromophenol blue) and heating at 80 °C for 10 min. For detection of CD9 and CD63, β-mercaptoethanol was omitted from the buffer. Protein samples (2 µg per well) were resolved on 4–20% Mini-PROTEAN® TGX™ Precast Protein Gels (Bio-Rad).

Following electrophoresis, proteins were transferred onto 0.45 µm nitrocellulose membranes (Amersham Protran) and blocked in 5% non-fat milk in PBS for 1 h at room temperature. Membranes were then incubated overnight at 4 °C with gentle shaking in primary antibodies diluted 1:1000 in 3% non-fat milk in PBS: anti-CD9 (clone EPR23105-125, cat. no. ab263019), anti-CD63 (clone EPR5702, cat. no. ab1340045), anti-TSG101 (clone EPR7130(B), cat. no. ab125011), and anti-HSP70 (clone EPR16892, cat. no. ab181606) (all from Abcam), or anti-ApoA1 (clone B-10, cat. no. sc-376818) (Santa Cruz Biotechnology).

Following primary antibody incubation, membranes were washed twice with PBS-T (0.05% Tween-20 in PBS) and once with PBS, then incubated for 1 h at room temperature with HRP-conjugated goat anti-rabbit or anti-mouse IgG secondary antibodies (Jackson ImmunoResearch, West Grove, PA, USA) diluted 1:3000 in 3% non-fat milk in PBS. Blots were washed as described above and developed using Clarity Max Western ECL substrate (Bio-Rad) and imaged with a ChemiDoc Imaging System (Bio-Rad).

### Exo-insert proteomic sEV preparation

Single-pot proteomic sample preparation of sEVs was performed using novel Exo-Insert workflow, enabling lysis, digestion, and LC–MS sample preparation without intermediate transfer steps in a single glass chromatographic insert. An equivalent volume of isolated Ti-EVs and CCM-EVs corresponding to 0.5, 1, 2, 4 µg (in triplicates) of protein was diluted in 3 ml PBS and ultracentrifuged at 100,000 × gmax (42□900 rpm) for 67 min at 4°C using a TLA-110 rotor (Optima MAX-XP Ultracentrifuge, Beckman Coulter) to pellet sEVs and remove residual compounds potentially incompatible with mass spectrometry. The pellets were dissolved in 20 µl of lysis and solubilization buffer composed from 0.2% (w/v) n-dodecyl-β-D-maltoside (DDM) and 10 mM dithiothreitol (DTT) in LC–MS-grade water freshly prepared prior to use. The sEV pellets obtained by ultracentrifugation were solubilized directly in the original ultracentrifugation tubes. The pellet location was identified based on prior tube marking and 20 µl of DDM lysis buffer were added to the pellet. Pellets were thoroughly resuspended by repeated pipetting and gentle scraping to maximize recovery. The resulting suspension was transferred directly into a 200 µl glass chromatographic insert with a conical bottom. Samples underwent two snap-freeze cycles in liquid nitrogen followed by sonification in a water bath for 5 min and incubation on ice for 30 min to facilitate complete lysis and protein extraction. Alkylation was performed by adding 5 µL of 50 mM iodoacetamide (IAA), resulting in a final concentration of approximately 10 mM. Samples were incubated in the dark to prevent reagent degradation with inserts covered to minimize evaporation. To reduce the DDM concentration to approximately 0.02%, 114 µL of 25 mM ammonium bicarbonate was added and samples were briefly vortexed. Proteins were digested by adding 0.4 µg sequencing-grade trypsin (Promega) to each insert. Samples were again vortexed, sealed, and incubated overnight at 37 °C. Following digestion, samples evaporated to dryness using a SpeedVac concentrator, taking care to avoid over-drying. Dried peptides were reconstituted in 15 µL of loading buffer consisting of 2.5% *(v/v)* acetonitrile and 0.08% *(v/v)* trifluoroacetic acid in LC–MS-grade water. Samples were transferred in the inserts into chromatographic autosampler vials and sealed by caps prior to LC–MS analysis. During mass spectrometric analysis, 7 µL of each sample was injected for data-dependent acquisition (DDA) and 7 µL for data-independent acquisition (DIA).

### Exo-SP3 sEV sample preparation

Three replicates of Ti-EVs and CCM-EVs suspensions containing equivalent 0.5, 1, 2, 4 µg of protein were brought up to 40 µl with 4% SDS in 100 mM ammonium bicarbonate, resulting in ∼2% SDS. Then, samples were reduced with 10 mM DTT for 30 min at room temperature and alkylated with 20 mM iodoacetamide for 30 min at room temperature in darkness. 2 µl of pre-washed 1:1 mix of e3 and e7 SERA-MAG beads were added to the samples. Then 50 µl of absolute ethanol was added, and samples were vortexed for 15 min. Next, beads were washed 4 times with 300 µl of 80% ethanol for 10 min with vortexing. After last wash, beads were dried under vacuum for 15 min and resuspended in 40 µl of 100 mM ammonium bicarbonate. Tryptic digestion was initiated by adding 1 µg of trypsin to each sample, and was carried out overnight at 37°C. Then, supernatant was separated from the beads and peptides were desalted with identical bead mix as used before. 1 µl of mix was added to peptide solution, and then 1 ml of pure acetonitrile. Samples were vortexed for 20 min, and beads were washed once with 0.5 ml of pure acetonitrile for 5 min with vortexing. Then, beads were dried under vacuum and peptides were eluted with 30 µl of 2% DMSO in water for 30 min. TFA concentration was set to 0.1 % prior LC-MS/MS

### SDS-FASP (FASP_SDS) method lysis

Three replicates of Ti-EVs containing 1, 4, 15 µg equivalent of protein were lysed in 50 µl of 4% (w/v) SDS, 100 mM Tris-HCl (pH 7.6), and 0.1 M DTT as described in Wu et al. [10]. Samples were incubated 30 min on ice to ensure complete solubilization and reduction of proteins.

### Urea-FASP (FASP_UA) method lysis

Three replicates of CCM-EVs containing 1, 4, 15 µg equivalent of protein in 17.4 µl PBS were diluted 4.6-fold with 8 M urea, 0.1 M Tris/HCl pH 8.5 buffer (UA buffer). The diluted samples were subjected to two cycles of snap-freezing in liquid nitrogen and subsequently stored at −80 °C overnight. After thawing, samples were centrifuged at 15,000 × g for 15 min at 9 °C to isolate the cleared supernatants.

### Filter-Aided Sample Preparation (FASP)

Protein digestion in urea lysed and SDS lysed supernatants was performed using the modified filter-aided sample preparation (FASP) method inspired by Wiśniewski et al. [16]. The protein lysates were loaded onto 10 kDa molecular weight cut-off centrifugal filter units (Microcon, Ultracel PL-10, 10 kDa) followed by UA buffer addition up to 200 µL of total volume. The unit was centrifuged at 14000 × g for 15 min at 20 °C. For reduction, 100 µL of UA buffer and 20 µL of tris(2-carboxyethyl) phosphine (TCEP) solution were added to the filter unit. The contents were mixed gently by pipetting and incubated for 30 min at 37 °C with shaking at 600 rpm in a thermomixer. After incubation, the filter unit was centrifuged at 14000 × g for 15 min at 20 °C. Alkylation was performed by adding 100 µL of UA buffer and 20 µL of 300 mM iodoacetamide to the filter. The solution was mixed gently and incubated for 20 min at room temperature in the dark with shaking at 600 rpm. The filter was subsequently centrifuged at 14000 × g for 15 min at 20 °C.

To remove urea and excess reagents, buffer exchange was performed using additional three washes with 100 µl of UA buffer followed by three washes using 100 µl of 100 mM ammonium bicarbonate each wash followed by centrifugation at 14000 × g for 20 min at 20 °C. After buffer exchange, the collection tube was replaced with a fresh tube. Trypsin was added at an enzyme-to-protein ratio of approximately 1:30 *(w/w)* in 100 µl of 50 mM ammonium bicarbonate. Samples were incubated overnight at 37 °C with shaking in a thermomixer.

### FASP peptide cleanup using C18 desalting columns

Peptide desalting was performed using Micro SpinColumns packed with C18 reversed-phase material (Harvard Apparatus, USA) according to a protocol inspired by Bouchal et al. [48].

Prior to sample loading, the C18 columns were activated with 200 µl of acetonitrile (ACN) containing 0.1% *(v/v)* formic acid (FA), followed by centrifugation at 100 × g for 3 min. This activation step was performed twice. After activation, the columns were equilibrated with 200 µl of water containing 0.1% *(v/v)* FA and centrifuged at 300 × g for 3 min. Subsequently, an additional 200 µl of water with 0.1% FA was added, and the column was allowed to hydrate for 15 min at room temperature. The column was then centrifuged again at 300 × g for 3 min.

The collection tube was emptied, and the peptide sample was loaded onto the equilibrated C18 column. The column was centrifuged at 500 × g for 3 min to allow peptide binding to the stationary phase. Bound peptides were washed three times with 200 µl of water containing 0.1% *(v/v)* FA. Each wash was followed by centrifugation at 500 × g for 3 min. After washing, the collection tube was replaced with a fresh tube, and peptides were sequentially eluted in three steps. First, 200 µl of 50% *(v/v)* ACN with 0.1% *(v/v)* formic acid was added followed by centrifugation at 500 × g for 3 min. Then, 200 µl of 80% *(v/v)* ACN with 0.1% *(v/v)* FA was added to the spin column followed by centrifugation at 500 × g for 3 min. The third elution step was done by 200 µl of 100% ACN with 0.1% *(v/v)* FA followed by centrifugation at 500 × g for 3 min. The eluates were combined and evaporated to dryness using a SpeedVac concentrator. Dried peptides were stored at −20°C until further analysis. Prior analysis tryptic peptides were dissolved in 15 µl of 2.5% ACN, 0.08% trifluoroacetic acid (TFA) in LC-MS grade water.

### LC–MS/MS analysis (data-dependent acquisition)

The volume of each injection was set to 7 µl. Peptide separation was performed using a trap-and-elute nanoLC setup consisting of a loading pump and a nano pump with a 90 min separation method. The loading pump operated under isocratic conditions at a flow rate of 5 µl/min using a loading buffer composed of 2.5% ACN and 0.08% TFA in LC-MS grade water. Analytical separation was achieved on the nanoLC using a linear gradient at a flow rate of 300 nl/min. Mobile phase A consisted of 0.1% formic acid (FA) in LC-MS grade water, while mobile phase B contained 80% ACN with 0.1% FA. The gradient profile was as follows: 2.5% B for 10 min, increased linearly to 40% B over 60 min, then to 99% B over 2 min, held at 99% B for 8 min, decreased to 2.5% B over 2 min, followed by column re-equilibration at 2.5% B for 8 min. Mass spectrometric analysis was performed in data-dependent acquisition (DDA) mode. Full MS scans were acquired over an m/z range of 350–1200 at a resolution of 120,000 with a normalized AGC target of 300%, automatic maximum injection time, one microscan, and profile-mode acquisition. MS/MS spectra were acquired at a resolution of 15,000 using an isolation window of 2 m/z, a maximum injection time of 40 msec, and one microscan. Data were recorded in centroid mode with normalized collision energy applied. Precursor ions were selected based on monoisotopic peak determination for peptides, with an intensity threshold of 5 × 10^3^ and charge states restricted to 2+ to 6+. For each full MS scan, the top 15 most intense precursor ions were selected for fragmentation. Dynamic exclusion was set to 20 sec to avoid repeated sequencing of the same precursor ions.

### LC–MS/MS analysis (data-independent acquisition)

Data-independent acquisition (DIA) measurements were performed using the same chromatographic conditions as described above for the DDA method. The DIA method consisted of a full MS scan followed by DIA MS/MS scans. Full MS scans were acquired over an m/z range of 350–1450 at a resolution of 60,000 with a normalized AGC target of 300%. The maximum injection time was set to 100 msec, with one microscan, and data were acquired in profile mode. DIA MS/MS scans were acquired across a precursor m/z range of 350–1100 using isolation windows of 12 m/z with a 1 m/z overlap. Fragmentation was performed using higher-energy collisional dissociation (HCD) with a normalized collision energy of 30%. MS/MS spectra were acquired at a resolution of 30,000 (at m/z 200), with an RF lens setting of 50% and a normalized AGC target of 1000%. The maximum injection time was set to automatic, with one microscan, and data were acquired in profile mode.

### Spectral library generation and database search

DIA data were processed in PEAKS Studio for spectral library generation using a human Swiss-Prot and TrEmbl protein database (including isoforms) supplemented with a common contaminant database. Protein identifications sharing the same set of identified peptides were grouped into protein groups for downstream analyses. Searches were performed with a precursor mass tolerance of 10 ppm and a fragment mass tolerance of 0.1 Da, assuming trypsin specificity with up to one missed cleavage. Peptides ranging from 6 to 45 amino acids in length were considered. Carbamidomethylation was set as a fixed modification, while protein N-terminal acetylation and methionine oxidation were included as variable modifications, with a maximum of two variable modifications allowed per peptide. Deep learning–based scoring enhancement was enabled. Results were filtered to a 1% false discovery rate (FDR) at both peptide-spectrum match (PSM) and protein group levels, and de novo sequencing results with an ALC score ≥50% were retained. De novo searches were conducted using the same mass tolerances, enzyme specificity, and modification settings as described for the database search. Identification of each protein group from DDA data was accompanied by spectral counting across all methods and replicates.

### DIA data processing and label-free quantification

For protein quantification, DIA data were analyzed using the previously generated spectral library, which was further expanded with identifications derived directly from the DIA datasets. Searches were conducted against the same human Swiss-Prot and TrEMBL protein database (including isoforms) supplemented with a common contaminant database. Search parameters, including mass tolerances, enzyme specificity, peptide length range, and modification settings, were kept consistent with those used for spectral library generation.

Label-free quantification was performed using precursor ion features with charge states ranging from 1+ to 5+, without additional filtering based on feature quality or intensity. Protein inference included modified proteins, and quantification was based on at least one peptide per protein. Quantitative values were calculated using the MaxLFQ algorithm, without fold-change cut-offs applied during quantification. Peptide features were aligned across samples using retention time correction to ensure complete peak integration. Signal intensities were subsequently normalized by total ion current (TIC).

### Data visualisation

Extracted values from both DIA and DDA data were processed and visualised in R programming language using ggplot2 version 3.5.2 and pheatmap version 1.0.13. Umap was created in R programming language using uwot package version 0.2.4 and PCA in PCAtools version 2.18.0.

## ETHICS APPROVAL AND PATIENT CONSENT

The study was conducted with ethical approval from the Independent Bioethics Committee for Scientific Research at the Medical University of Gdańsk (approval no. NKBBN/880-329/2016). All patients provided written informed consent prior to sample collection.

## AUTHOR CONTRIBUTIONS

J.F.: Conceptualization, Methodology, Data analysis, Formal analysis, Visualization, Data curation, Data interpretation, Drafting, Review & Editing.

A.P.: Conceptualization, Methodology, Investigation, Formal analysis, Review & Editing, Drafting.

A.B.: Conceptualization, Methodology, Drafting, Formal analysis, Review & Editing, Visualization.

I.P., B.S., S.B, T.M., W.R.: Resources, Review & Editing.

N.M.T.: Resources, Funding acquisition; Review & Editing.

S.K.: Conceptualization, Supervision, Resources, Funding acquisition, Review & Editing.

