## Supplementary figure_1 for "Benchmarking five extracellular vesicle proteomics workflows, including low-input Exo-insert and Exo-SP3, for deep mass spectrometry profiling"

1 ug sEV – FASP_SDS

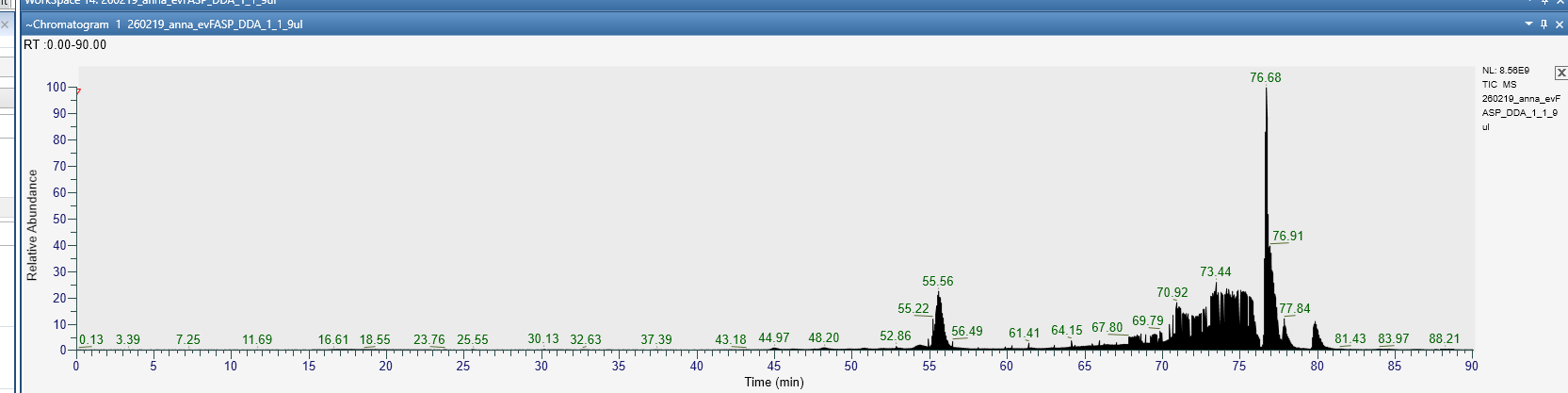

4 ug sEV – FASP_SDS

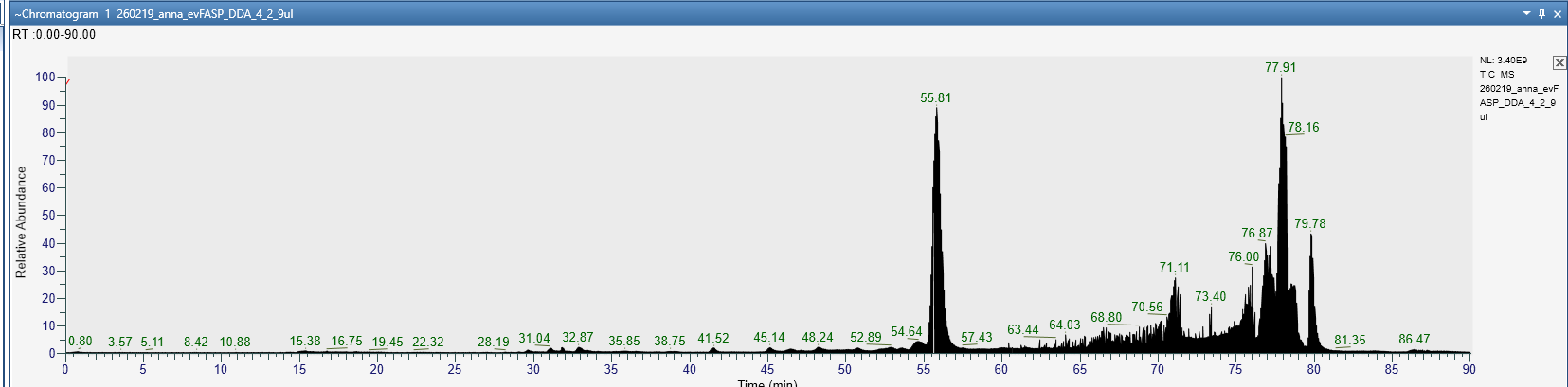

15 ug sEV – FASP_SDS

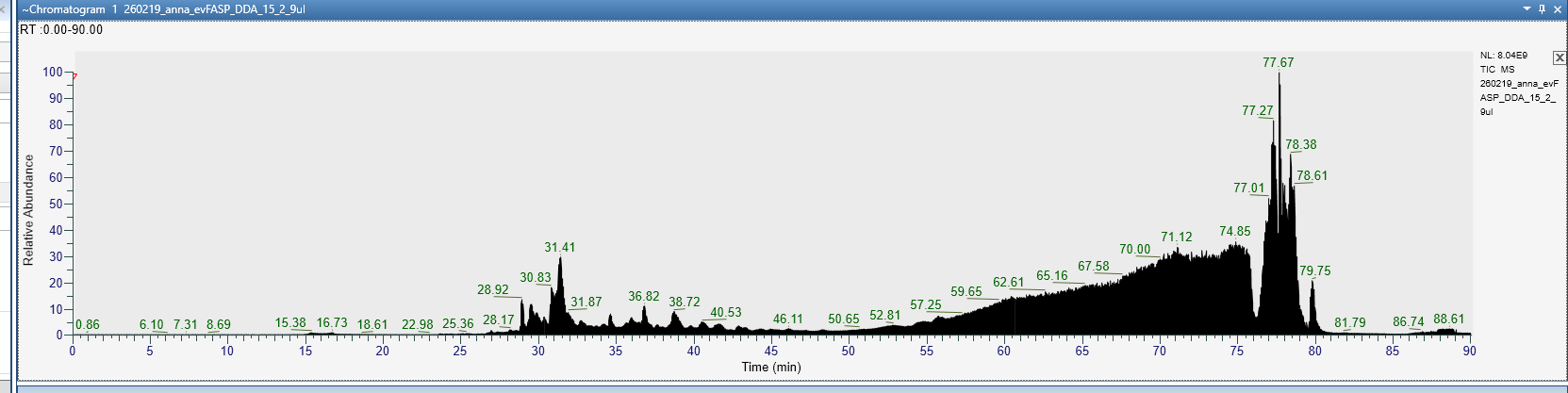

***Supplementary Figure 1.*** *Total ion chromatograms (TICs) of single replicates of 1, 4, and 15 µg Ti-EVs processed by SDS-based filter-aided sample preparation (SDS-FASP) and injected as a single LC-MS/MS run. The TICs indicate uneven peptide separation and compression of elution toward the end of the chromatograms. This pattern may reflect the atypical single-injection setup required by the limited sEV input, in contrast to canonical workflows where the resulting digest is typically divided across multiple injections. As a consequence, residual SDS and other matrix components that would otherwise be distributed across several runs may have been concentrated into one injection, potentially impairing chromatographic separation.*
