## Supplementary figure_2 for "Benchmarking five extracellular vesicle proteomics workflows, including low-input Exo-insert and Exo-SP3, for deep mass spectrometry profiling"

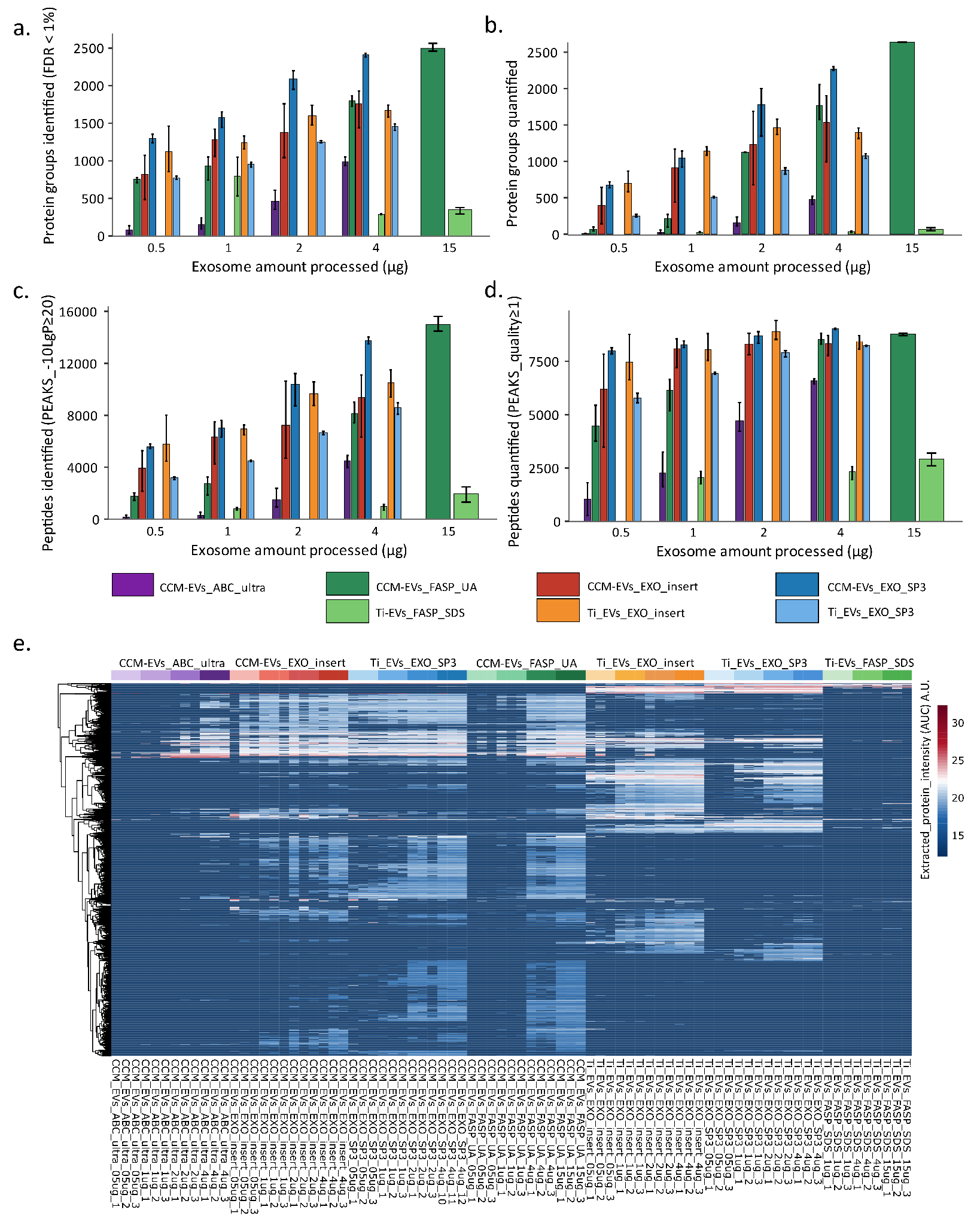

***Supplementary figure 2.*** *Comprehensive comparison of sEV proteome coverage across all preparation methods, input amounts, and acquisition strategies. CCM-EVs and Ti-EVs were processed using five preparation methods (Exo-insert, Exo-SP3, ABC, FASP_UA, and FASP_SDS) across multiple sEV input amounts (0.5–4 µg; up to 15 µg for FASP methods), each analyzed in technical triplicate. Due to sample availability constraints, FASP_UA and ABC_ultra were applied only to CCM-EVS, while FASP_SDS was applied only to Ti-EVs. Peptide samples were split for analysis by DDA and DIA on an Exploris 480 platform, and datasets were processed separately. (a) Stacked bar plots showing the mean number of protein groups identified by DDA. (b) Stacked bar plots showing the mean number of protein groups quantified by DIA. (c) Stacked bar plots showing the mean number of peptides identified by DDA. (d) Stacked bar plots showing the mean number of peptides quantified by DIA. Error bars in bar plots (a-d) show minimum and maximum value. (e) Heatmap of DIA-derived protein intensities across all conditions.*
